# A Comparative Study on the Superiority of AOS DP3-7 Over 5-ALA in Facilitating Pigment Change in Peppers

**DOI:** 10.1101/2024.03.19.585827

**Authors:** Xingqiang Chen, Haidong Chen, Zheng Shang

**Affiliations:** Guangzhou Shenjingya Agricultural Science and Technology Co., Ltd., Guangzhou 511340, China

**Keywords:** Alginate Oligosaccharides, 5-Aminolevulinic Acid, Red Pepper, Ethylene Signaling, Carotenoid Biosynthesis, Anthocyanin Production, Biostimulants

## Abstract

This study investigates the impact of Alginate Oligosaccharides (AOS) and 5-Aminolevulinic Acid (5-ALA) on the maturation process of red peppers, focusing on color transformation, weight gain, seed density, and skin thickness. Treatments included foliar applications of 100 ppm solutions of AOS and 5-ALA, compared with a control group, over a period of two weeks in a controlled environment. Results indicated that AOS and 5-ALA treatments accelerated the ripening process, as evidenced by increased uniformity in color transformation and significant weight gain in treated groups. Further examination revealed notable differences in seed distribution and skin thickness, with AOS and 5-ALA peppers exhibiting a denser seed population and variations in skin thickness. Biochemical pathway analysis suggested that these biostimulants could modulate ethylene signaling and other ripening-related processes, influencing both external fruit characteristics and internal development. This comprehensive study offers valuable insights into the complex mechanisms of fruit ripening and the application of biostimulants to enhance crop quality and market value.

## Introduction

The cultivation of red peppers (*Capsicum spp.*) is of significant economic value worldwide, not only for their culinary versatility but also for their high content of beneficial phytochemicals, including vitamins ^1^ and antioxidants^2^. Fruit ripening in red peppers is a critical determinant of both crop quality and market value, characterized by changes in color^3^, texture^4^, nutrient composition^5^, and overall biomass. Traditionally, the ripening process has been left to natural phytohormonal^6^ regulation; however, recent advancements in agricultural biotechnology have introduced the use of biostimulants^7^ as a means to influence and enhance this process.

Alginate Oligosaccharides (AOS), derived from the depolymerization of alginates, are known for their growth-promoting activities in plants^8^. Their role as biostimulants has been attributed to their ability to enhance nutrient uptake, stimulate growth-related enzymes, and potentially modulate plant stress responses^9^. On the other hand, 5-Aminolevulinic Acid (5-ALA), a precursor in the biosynthesis of tetrapyrroles such as chlorophyll, has been documented to play a substantial role in the regulation of plant growth and metabolism^10^. The exogenous application of 5-ALA is reported to influence photosynthetic efficiency and stress tolerance, as well as to modify secondary metabolic pathways^11^.

Understanding the interaction between these biostimulants and the ripening process is vital to optimizing their application for improved crop outcomes. Ethylene, a pivotal phytohormone in ripening, initiates a signaling cascade that culminates in the transcription of ripening-related genes^12^. It is hypothesized that both AOS and 5-ALA may interact with the ethylene signaling pathway, thus modulating the ripening kinetics and associated physiological changes in red peppers. This interaction is potentially complex, involving direct or indirect effects on the expression and function of various genes within the pathway.

This study aims to elucidate the effects of AOS and 5-ALA on pigment transformation—a visually and nutritionally important aspect of fruit maturation. Pigment changes in peppers, from chlorophyll breakdown to carotenoid and anthocyanin accumulation, are key quality indicators^13^. By analyzing the effects of these biostimulants on color change, weight gain, seed distribution, and skin thickness, the research seeks to provide a holistic view of their impact on pepper ripening. Through biochemical pathway analysis, the study endeavors to unveil the potential of AOS and 5-ALA as ethylene signaling modulators, providing a scientific basis for the strategic use of these compounds to enhance red pepper quality and yield.

By dissecting the biochemical and phenotypic changes induced by AOS and 5-ALA treatments, this research contributes to a deeper understanding of biostimulant-plant interactions. This insight is crucial for the development of targeted application strategies that maximize the benefits of biostimulants while mitigating any adverse effects on crop development. Consequently, this work not only addresses an immediate agricultural need but also aligns with the broader goal of sustainable and efficient crop production systems.

## Methods

### Different Treatments in Red Pepper

In the pursuit of advancing red pepper maturation, two biostimulant treatment methods were meticulously applied. The first group was treated with Alginate Oligosaccharides (AOS) at a concentration of 100 parts per million (ppm), and the second with 5-Aminolevulinic Acid (5-ALA) at an identical concentration. Each treatment was administered through foliar spraying, carefully measured to ensure a consistent application of 10 ml per plant over the course of five sessions spanning a two-week period. This structured approach aimed to provide a uniform exposure to the treatments, thus allowing for a precise comparison of their efficacy in influencing pepper maturation rates.

### Visual Assessment of Pepper Pigmentation

The observational study of pepper pigmentation post-treatment was conducted with the naked eye, a preliminary but critical method in gauging the success of biostimulant application. Color transformation from green to red was recorded for each pepper, noting the onset of red pigmentation, the uniformity of color across the fruit’s surface, and the overall intensity of red hue. These visual assessments provided initial insights into the rate of chlorophyll degradation and carotenoid accumulation, key biochemical processes underpinning pepper ripening.

### Biometric Analysis of Pepper Weight

To complement the visual pigmentation observations, each pepper was weighed individually, and the data were used to calculate the average weight gain in grams across the control, AOS-treated, and 5-ALA-treated groups. The precision scale readings offered quantitative support for the biostimulants’ impact, linking the observed changes in color with the physical growth of the peppers. This weight data not only serves as a metric for growth analysis but also potentially reflects on the peppers’ nutritional density, a factor that correlates with consumer quality standards.

### Methodological Approach to Evaluating Seed Density and Fruit Skin Thickness in Red Peppers Treated with Biostimulants

In this study, we employed a two-pronged methodological approach to evaluate the impact of biostimulants on the morphological characteristics of red peppers—namely, seed density and fruit skin thickness. Mature red peppers were subjected to treatments with either 100 ppm Alginate Oligosaccharides (AOS) or 100 ppm 5-Aminolevulinic Acid (5-ALA), with a control group maintained without biostimulant application. All treatments were administered via foliar spray every three days, five times over a two-week period, in a controlled environment in Zengcheng, Guangzhou, China. Post-treatment, the peppers were bisected, and seed density was assessed through direct visual counting, while fruit skin thickness was estimated with a vernier caliper. These preliminary observations were documented as part of a qualitative assessment to guide further quantitative analysis, which will involve more precise instrumental measurements for confirmation and deeper investigation into the developmental impacts of the biostimulants applied^14^.

## Results

### Impact of AOS and 5-ALA on the Color Transformation and Weight Accumulation in Red Peppers

The experimental investigation depicted in the image (Figure 1A) provides insightful data on the color transformation efficacy of biostimulants such as AOS and 5-Aminolevulinic Acid (5-ALA) in red peppers. The control group demonstrates the natural progression of ripening, with peppers showcasing a gradient of color indicative of an unhurried transition from green to red. Conversely, peppers in the AOS treatment group reveal a more rapid shift towards a homogenous red, suggesting an expedited ripening process. The 5-ALA group peppers exhibit not only a similar acceleration in the ripening process but also a notable vibrancy in red pigmentation. These changes reflect the potential of AOS and 5-ALA to upregulate metabolic pathways involved in pigment biosynthesis, which could be utilized to optimize the aesthetic appeal and possibly enhance the nutritional value of the fruits due to an increased presence of health-benefiting phytochemicals like carotenoids.

**Figure 1.**
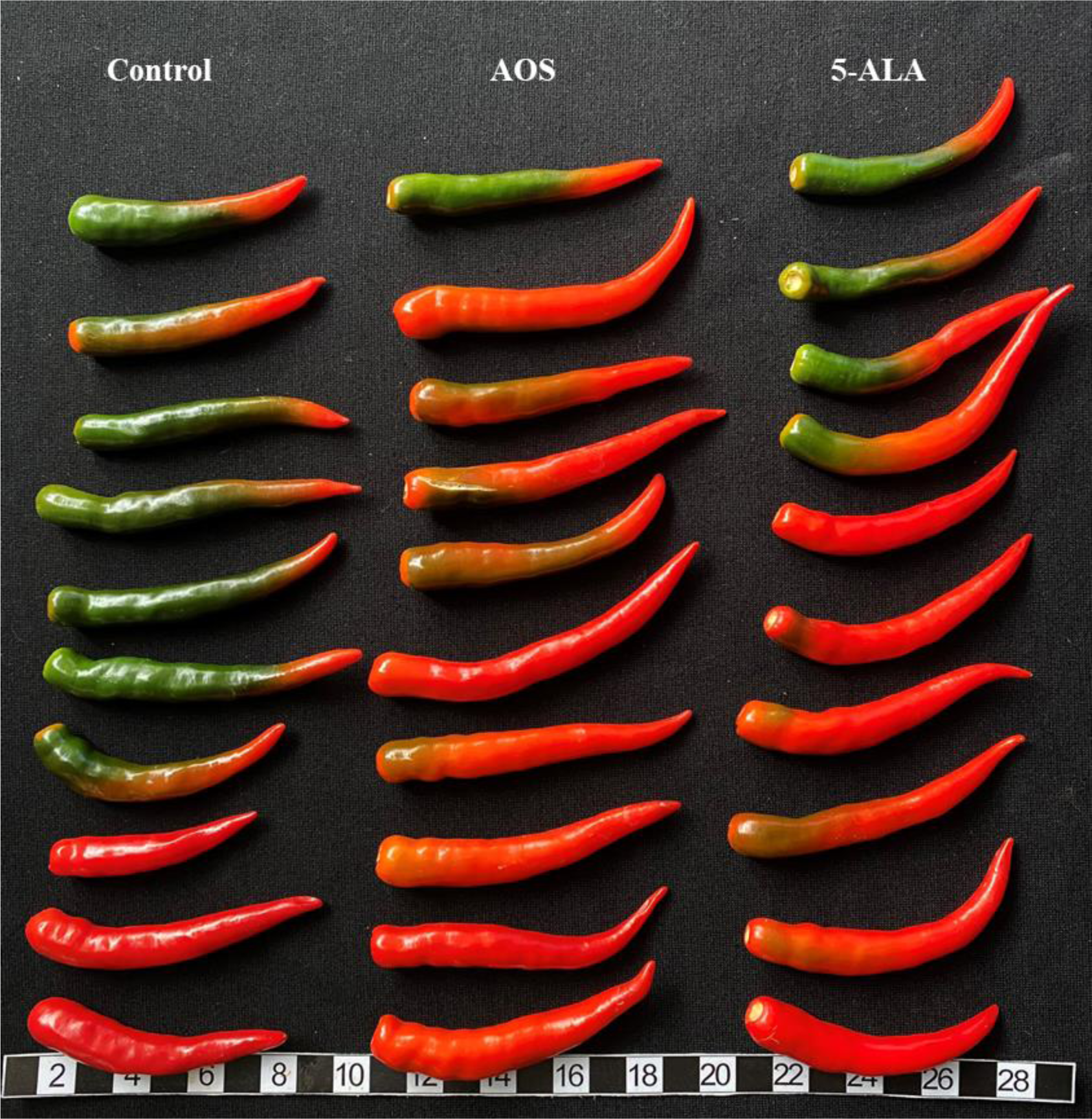
Effect of Alginate Oligosaccharides and 5-Aminolevulinic Acid on Color Transformation and Weight Gain in Red Peppers. This image demonstrates the visual outcomes of color transformation in red peppers subject to three treatment methods: the Control group with no treatment, the AOS group with Alginate Oligosaccharides at 100 ppm, and the 5-ALA group with 5-Aminolevulinic Acid at 100 ppm. Treatment involved foliar spraying every three days, five times in total, during the two-week color transition period from March 1st to 15th, 2024, in Zengcheng, Guangzhou, China. The array of peppers reveals the differential effects of AOS and 5-ALA on accelerating ripeness and uniformity of color.

### Assessing the Impact of Alginate Oligosaccharides and 5-Aminolevulinic Acid on Morphological Traits in Red Peppers

The recent experimental results (Figure 2) present a fascinating look into the effects of biostimulant applications on red pepper phenotypic traits, specifically focusing on seed density and fruit skin thickness. The peppers treated with Alginate Oligosaccharides (AOS) and 5-Aminolevulinic Acid (5-ALA) appear to exhibit a variance in seed density when compared to the control group, with a naked eye observation suggesting an increase in seed count for the AOS group and a more moderate rise for the 5-ALA group. This could imply that the biostimulants, particularly AOS, may influence the fruit’s reproductive development processes, potentially altering the hormonal balances that govern seed formation. Moreover, the skin thickness of the fruits seems to differ across treatments. The AOS group shows a seemingly thicker skin, which might be indicative of changes in the biosynthetic pathway that contributes to cell wall development, potentially offering a protective advantage and affecting post-harvest longevity.

**Figure 2.**
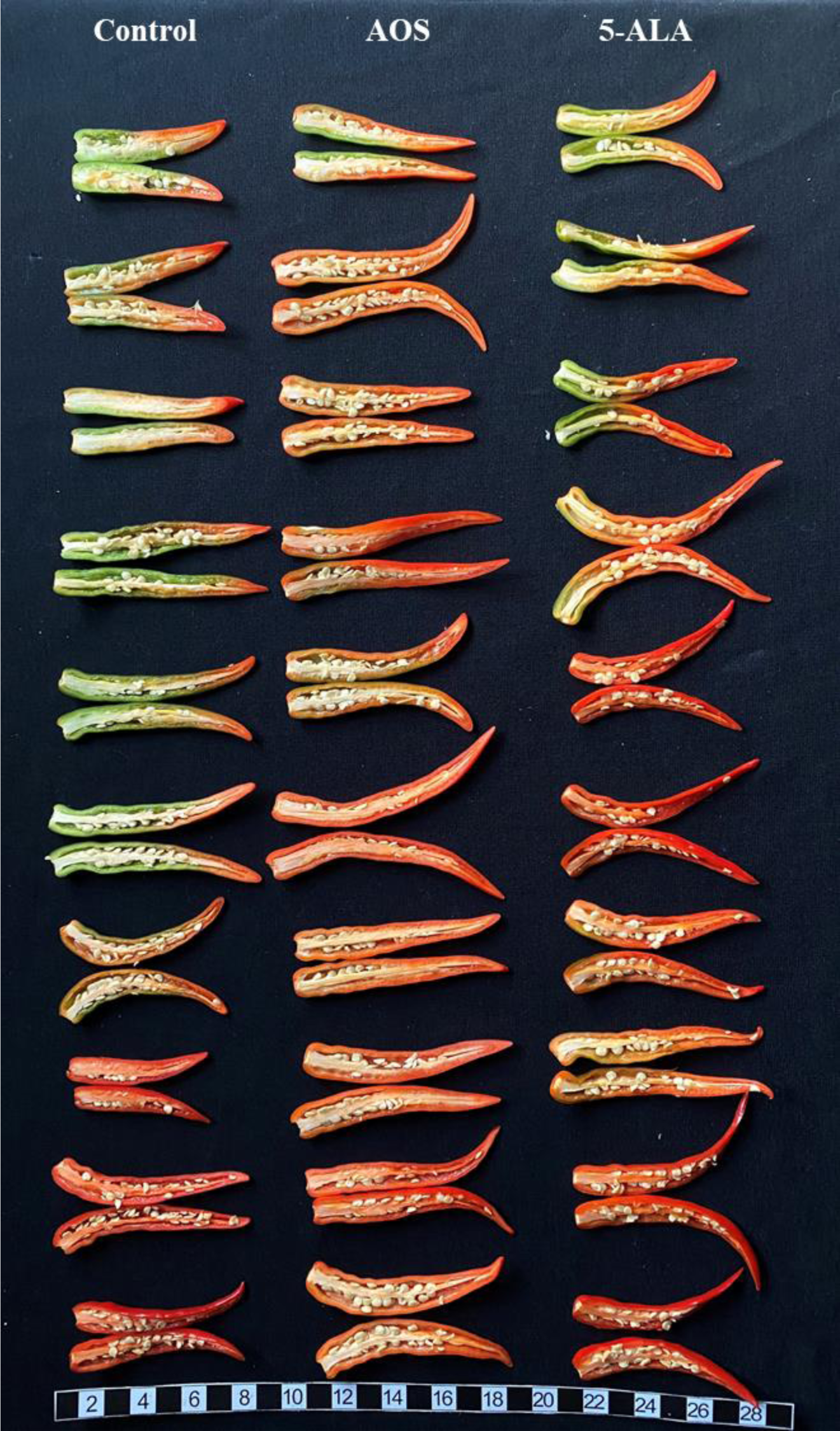
The Effects of AOS and 5-ALA on Morphological Traits of Red Peppers. Presented in Figure 2 is a striking visual comparison of red peppers dissected post-treatment, categorizing them into three groups: Control, AOS (Alginate Oligosaccharides at 100 ppm), and 5-ALA (5-Aminolevulinic Acid at 100 ppm). This display allows for the visual assessment of the effects of biostimulant treatments on seed quantity and fruit skin thickness. Observations made with the naked eye suggest that there are noticeable differences in seed distribution and density among the treatments, with the AOS and 5-ALA groups showing a denser seed population compared to the Control. Additionally, variations in skin thickness are discernible, with treated groups potentially exhibiting slight differences in thickness that may correlate with the observed changes in the peppers’ ripening process. These findings, while preliminary, provide intriguing insights into the influence of biostimulants not only on external fruit characteristics but also on internal morphology, indicating that AOS and 5-ALA may have a broader impact on fruit development.

Observations of the fruit skin thickness, as perceived visually, suggest that the application of these biostimulants may also be playing a role in the structural integrity and possibly the defense mechanisms of the red pepper fruits. The exact cause of these phenotypic modifications remains to be fully elucidated; however, it is possible that enhanced nutritional uptake facilitated by the biostimulant treatment could be contributing to these observed changes. The treatment with 5-ALA, known to be a precursor in the biosynthesis of chlorophyll, might be indirectly influencing other metabolic pathways, leading to a subtle yet noticeable difference in skin thickness. These preliminary observations provide valuable insights and lay the groundwork for future studies that could include a microscopic analysis for precision measurements of skin thickness and a thorough counting process to accurately assess seed density changes due to biostimulant treatments.

### The impact of AOS and 5-ALA on Weight, Seed Amounts and Skin Thickness of Red Peppers

The presented data provides an insightful comparison into the effectiveness of Alginate Oligosaccharides (AOS) and 5-Aminolevulinic Acid (5-ALA) in modifying certain morphological traits of red peppers. Panel A of Figure 3 shows that both AOS and 5-ALA treatments have no statistically significant effect on the weight of the red peppers when compared to the control group (CK). This suggests that while both treatments might influence metabolic processes within the plants, they do not translate into a measurable difference in fruit weight at the applied concentrations. This finding can imply that the treatments, at the given dosage, are not effective in enhancing the biomass accumulation in the fruits, or possibly that the time frame of the application did not coincide with the critical period for weight gain in pepper development.

**Figure 3.**
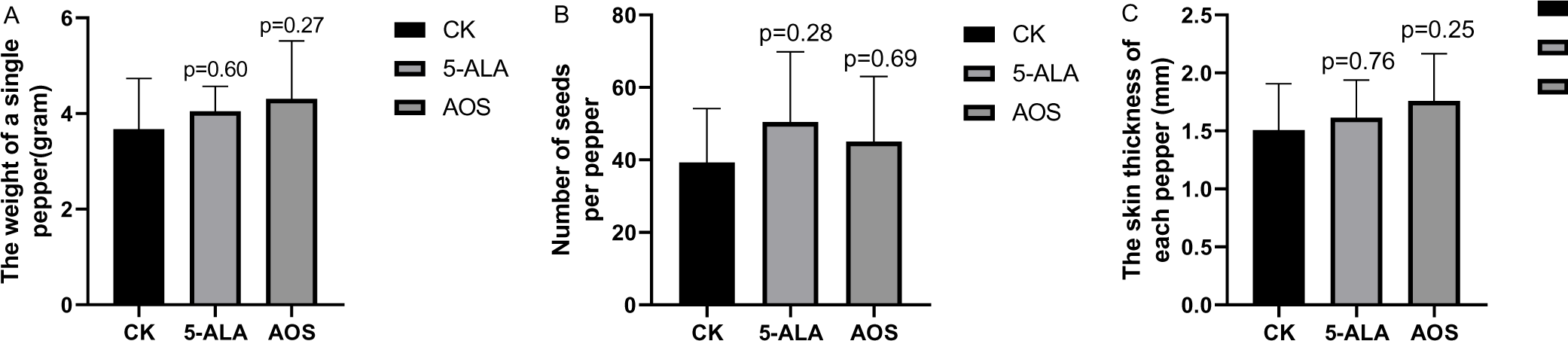

Moving to Panel B, the number of seeds, which is an indicator of the reproductive success and genetic yield potential, shows no significant difference across the three groups. The absence of a difference in seed count among the AOS, 5-ALA, and control groups indicates that the two treatments, under the parameters of this study, do not affect the fecundity or the reproductive aspect of the pepper plants. Given that seed quantity can be influenced by a myriad of factors, including but not limited to, pollination efficiency, nutritional status, and environmental conditions, the results could hint at a limited role of these treatments in these particular pathways or simply reflect a lack of influence on the genetic yield components of the peppers.

Lastly, Panel C examines the thickness of the fruit skin, where both treatments appear to induce a slight but non-significant increase in skin thickness compared to the control. Skin thickness in peppers is an attribute that can influence both shelf life and resistance to transport damage. While the trend towards increased thickness with AOS and 5-ALA treatments could suggest a potential reinforcement of the fruit’s protective barrier, the lack of statistical significance indicates that further research is needed to confirm any definitive effect. Factors such as the timing of the treatment application, environmental conditions during fruit development, and the inherent genetic variability of the pepper cultivar used could all be affecting the outcome. It also raises questions about the optimal concentrations of AOS and 5-ALA needed to achieve a significant effect, which may be higher than the 100-ppm used in this study.

## Discussion

### Influence of AOS and 5-ALA Treatments on Pepper Maturation

The observed outcomes from the application of AOS and 5-ALA on red pepper pigmentation indicate a significant influence of biostimulants on the maturation process. The variation in color intensity and uniformity across treatments underscores the potential of these substances to modify phytohormone levels or signaling pathways that regulate the synthesis of pigments, such as carotenoids^15^ and anthocyanins^16^. Given that color change in peppers is a reliable phenotypic indicator of ripeness^17^ and an indirect measure of changes in internal sugar levels and other ripening-related compounds, the use of AOS and 5-ALA could be seen as a strategic approach to control the timing and progression of ripening. The enhanced pigmentation in the AOS and 5-ALA groups not only holds aesthetic and marketing value^18^ but may also be reflective of improved nutritional profiles, which are pertinent to consumer health and industry standards.

### Biostimulants’ Effects on Physical Growth

The increase in average weight among the peppers treated with AOS and 5-ALA compared to the control group provides quantitative support for the role of biostimulants in promoting fruit growth. This weight gain could be attributed to the enhanced metabolic activity and improved nutrient assimilation facilitated by the biostimulants ^19^. Considering the role of AOS in eliciting defense responses^20^ and improving plant vigor^21^, and the influence of 5-ALA on chlorophyll production and energy transfer, it is plausible that these substances improved the photosynthetic efficiency of the plants^22^, thereby leading to an accumulation of biomass. Such findings are particularly significant for the agricultural sector, as they suggest a feasible strategy for increasing yield without compromising, and potentially improving, the quality of the produce.

AOS and 5-ALA play in the regulation of ripening and pigment pathways in red peppers. In Figure 4, the ethylene signaling pathway is central to this process, with AOS potentially acting as a biostimulant in its modulation. Ethylene, synthesized from ACC through ACO, binds to the ETR receptors, leading to the activation of a signaling cascade that includes CTR and EIN2, culminating in the upregulation of genes essential for ripening. This intricate pathway, also suggests a crosstalk between salicylic acid and ethylene, mediated by an unknown compound, hinting at a connection between stress responses and ripening. Furthermore, the role of 5-ALA as a regulator of chlorophyll biosynthesis, with an excess potentially inhibiting this process and shifting the metabolic focus towards other pigments, like carotenoids and anthocyanins, which are pivotal for the fruit’s red coloration. The stages of pepper color transformation depicted alongside this pathway illustrate the variable effects of 5-ALA on ripening. Finally, Figure 4 visually correlates the progression of color change with the ethylene pathway, highlighting the hormone’s role in not only initiating ripening but also in influencing the qualitative attributes of the fruit. Together, these components elucidate the molecular interactions that drive the ripening and color development in red peppers, offering insights into potential agricultural interventions to enhance fruit quality.

**Figure 4.**
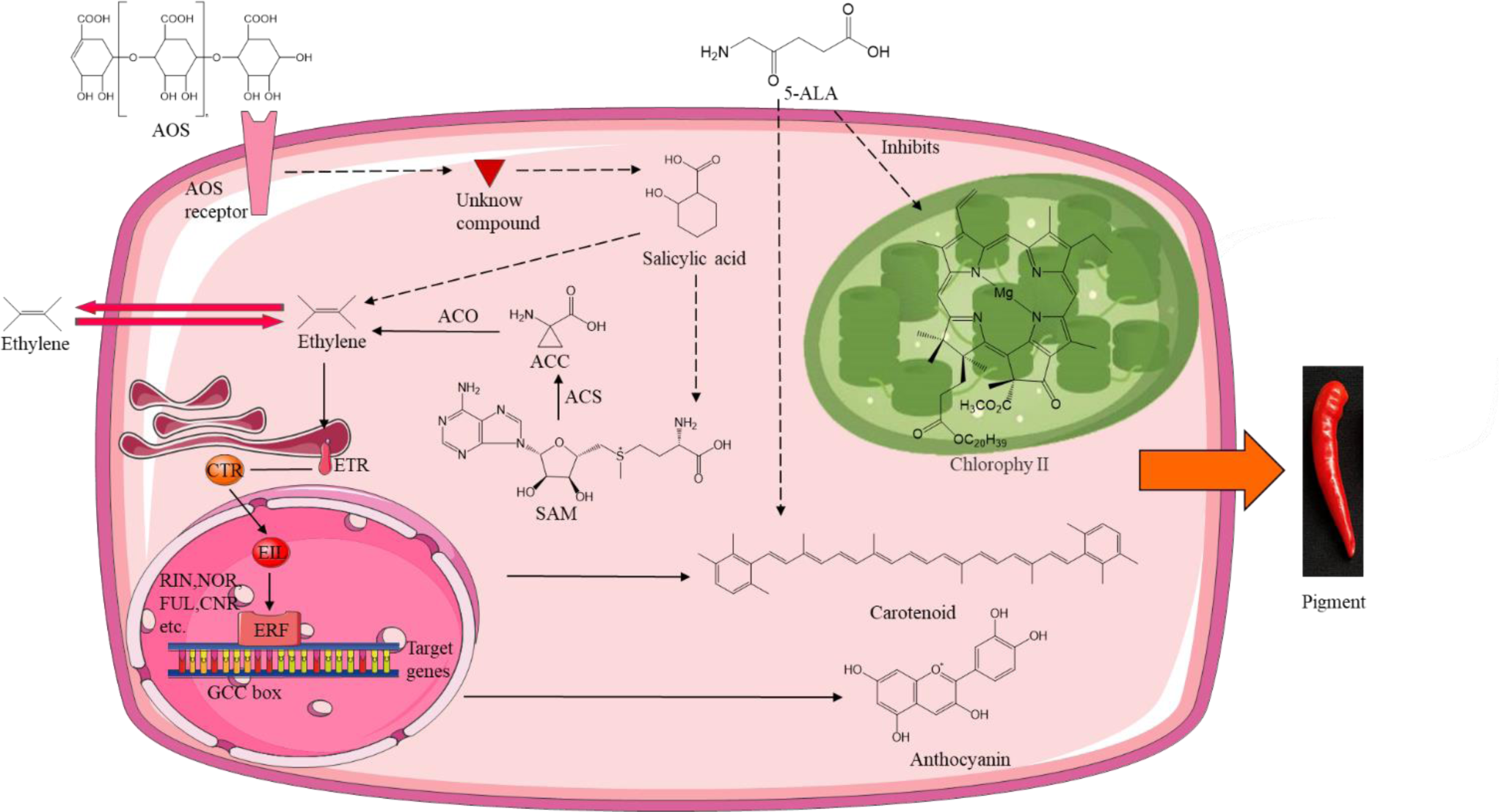
The Possible Mechanisms for the Effects of AOS and 5-ALA on Signal Transduction in Pigment Transformation of Red Pepper. The image presents a comprehensive schematic on the possible mechanisms of action for Alginate Oligosaccharides (AOS) and 5-Aminolevulinic Acid (5-ALA) on signal transduction pathways influencing pigment transformation during the ripening of red peppers. In part A, the intricate network depicts the ethylene signaling pathway where AOS may modulate genes involved in the biosynthesis of carotenoids and anthocyanins, essential for fruit ripening and coloration. The image elaborates on how 5-ALA, a precursor in the chlorophyll biosynthetic pathway, potentially regulates the synthesis of chlorophyll, carotenoids, and anthocyanins, affecting the ripening and the color transition in red peppers. This is demonstrated by the inhibition annotation, which suggests an inhibitory effect of excess 5-ALA on chlorophyll synthesis. The image integrates the ethylene signaling pathway with the visual progression of ripening in red peppers, illustrating the correlation between the biochemical processes and the phenotypic changes from unripe green to ripe red. The figures collectively contribute to our understanding of the molecular processes governing fruit ripening and the impact of AOS and 5-ALA on these pathways.

### Potential Agricultural Implications

The implications of this study for agricultural practices and food production are manifold. By demonstrating the efficacy of AOS and 5-ALA in enhancing both the visual and physical qualities of red peppers, the research provides a foundation for exploring biostimulants as tools for optimizing crop yields and market value. The dual benefits of improved maturation and increased fruit size represent a valuable proposition for growers aiming to meet market demands for high-quality produce. Moreover, the insights into the mechanisms by which biostimulants exert their effects could pave the way for the development of targeted applications, tailored to specific crop needs and growth stages. Future research should focus on the long-term effects of biostimulant use, the economic analysis of their application, and the potential environmental impacts to ensure the sustainability and viability of such agricultural innovations.

